# Single-cell Transcriptome Profiling of Post-treatment and Treatment-naïve Colorectal Cancer: Insights into Putative Mechanisms of Chemoresistance

**DOI:** 10.1101/2024.09.26.614712

**Authors:** Grigory A. Puzanov, Clémence Astier, Andrey A. Yurchenko, Gérôme Jules-Clement, Fabrice Andre, Aurélien Marabelle, Antoine Hollebecque, Sergey I. Nikolaev

## Abstract

Drug resistance remains a major clinical challenge in the treatment of colorectal cancer (CRC) with conventional chemotherapy. Analyzing changes within tumor cells and tumor microenvironment (TME) after treatment and in metastases is essential to understanding how resistance develops. In this study, we analyzed scRNA-seq data from 56 CRCs including treatment-naïve tumors and tumors treated with standard chemotherapy with the known response status (18 responders and 6 progressors). In our cohort, primary left-sided CRCs were associated with metastatic potential mesenchymal phenotype and with depleted B cells. In the post-treatment CRC, there was a high prevalence of dendritic cells (DC) in the TME in the response group. The DC-derived signature was associated with better survival in a large CRC cohort from the TCGA. In progressors there was an enrichment of pericyte-like fibroblasts, which appeared to be associated with poor survival in a CRC-TCGA cohort. Progressors also showed elevated fractions of exhausted CD8+ T memory cells suggesting a pro-inflammatory TME. In tumor cells of progressors group, we identified specific expression of chemo-protective markers *MTRNR2L1* and *CDX1*; and their co-expression with stemness-related immune-checkpoint *CD24*. In summary, scRNA-seq provides a valuable information for the discovery of prognostic markers, and reveals distinct features potentially underlying response to chemotherapy or disease progression in CRC.

## INTRODUCTION

Colorectal cancer (CRC) is the third most frequently diagnosed cancer type and the second by mortality rate, according to GLOBOCAN 2022 data[1]. The number of diagnosed cases of CRC world-wide in 2020 was 1.9 million, and it is expected to rise in future decades reaching 3.2 million by year 2040[2]. Along with surgery and radiotherapy, conventional chemotherapy, which usually includes 5-fluorouracile (5-FU), folinic acid and oxaliplatin or irinotecan, remains the main strategy for the treatment of CRC[3]. In some cases chemotherapy is used in combination with either anti-angiogenic or anti-EGFR treatments. Despite the fact that combined therapies improve the response rate, the prognosis of CRC remains poor, and the 5-year survival is still fewer than 20%[3, 4].

Pharmacologic resistance is the main cause of CRC recurrence and poor prognosis[5]. Resistance mechanisms of CRC cancer cells to cytotoxic effect of chemotherapy has been extensively studied. It was shown that population of cancer stem cells have an important multi-faceted role in chemo-resistance[6, 7]. Autophagy also can contribute to protection of tumor cells from the cytotoxic effects of chemotherapeutic agents[8]. A process of the drug-efflux from the cell by ATP transporters, which requires increased respiration, can be intensified in resistant tumors[9]. Some cell type signatures, such as derived from cancer-associated fibroblasts or immune cells, also showed association with survival in CRC[10–12]. Chronic inflammation has been described to lead to decreased effectiveness of chemotherapy and decreased survival[13]. More recently, evidence has emerged that prevention of apoptosis of resistant cells can cause inflammation in the tumor[14], thus promoting pro-tumoral immune microenvironment.

Tumor locations in left-sided or right-sided colon might also impact the response rate and mechanisms of resistance to treatment[15]. Right-sided CRC is known to better respond to immunotherapy[16], while left-sided CRC generally has a better prognosis and responds better to conventional chemotherapy[17]. Thereafter, there are numerous mechanisms in the CRC rendering a decreased sensitivity to chemotherapy, some of which are related to tumoral cells while others – to TME composition. In particular, recently it was shown on scRNA-seq that B cells are activated after preoperative chemotherapy and are reduced in liver metastases[18].

Since response to therapy in CRC depends on both, intrinsic features of tumor cells and on the TME, improvement of chemotherapy efficacy requires a better understanding of the interplay between cell types in the tumor[19, 20]. Recent scRNA-seq studies provided new insights in the complexity of TME and its interaction with the tumoral cells on large cancer type specific and pan-cancer cohorts[21–23]. These also allowed a detailed characterization of the subtypes of the tumor infiltrating cells and of the heterogeneous activity of transcriptional programs in cancer cells[21, 23].

In this study, using scRNA-seq data from a large CRC cohort, we identified major features of tumoral cells and TME, associated with treatment response status and other clinical characteristics such as metastatic status, tumor location and microsatellite instability (MSI). We demonstrated the association between a pro-inflammatory TME and progressive disease, and identified several cell type-specific gene signatures, which were correlated with response and outcome of CRC.

## RESULTS

### CRC cohort characteristics

Our total cohort included 32 untreated CRC (25 MSS and 7 MSI) and 24 post-treatment tumors including 18 responder and 6 progressors (Figure 1A). Within untreated cohort, the MSS sub-cohort included 19 left-sided and 6 right-sided CRCs (Figure 1A). This sub-cohort was enriched in low-grade tumors (stage I-II, 18/25) and half of it had a *KRAS* mutation (11/24). According to consensus molecular subtypes (CMSs) classification 3 tumors were CMS1, 4 – CMS2, 6 – CMS3 and 9 – CMS4 (Supplementary Fig. S1A). The majority of biopsies were taken from primary tumors (n=20), while only 5 were from metastases (Supplementary Table S1, Figure 1A). MSI sub-cohort included 5 right-sided and only 2 left-sided primary tumors.

**Figure 1.**
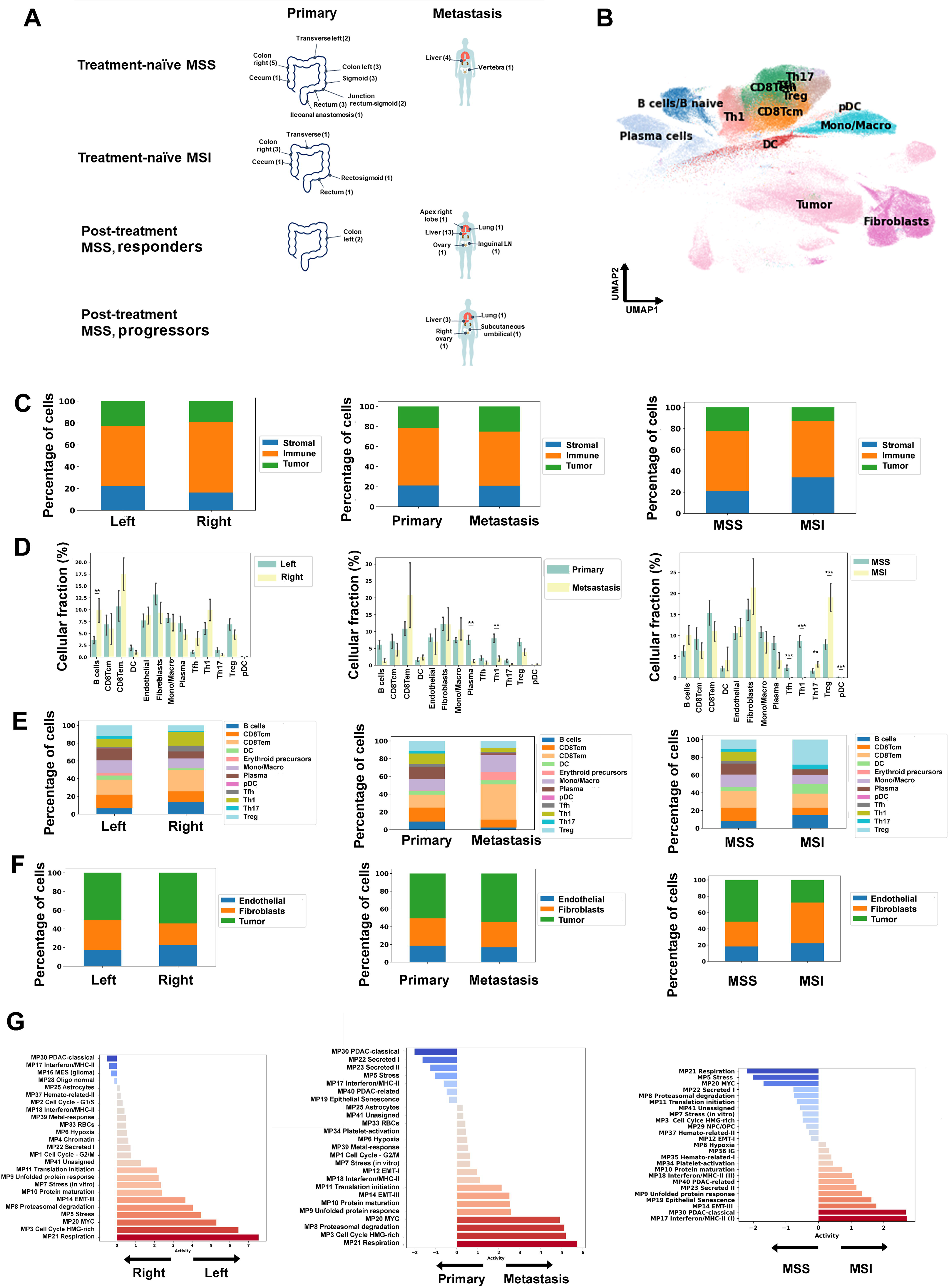
Characteristics of TME in treatment-naïve samples. **A**, Location of biopsies: primary and metastases. Orange arrows show comparisons made in the study: left and right MSS CRC, MSS and MSI CRC, primary and metastatic CRC, treatment-naïve MSS CRC and post-treatment MSS CRC, responders and progressors post-treatment MSS CRC. **B**, Uniform manifold approximation and projection (UMAP) embedding: high, 285,608 cells from 32 CRC treatment-naïve samples. **C**, TME proportions changes between left and right-sided tumors, between metastatic and primary tumors, between MSI high and MSI low tumors. Bars indicate cell percentages, whiskers indicate 0.75 confidence intervals, and asterisks indicate p-values for the Mann-Whitney test (* - p < 0.05, ** - p < 0.01, *** - p < 0.001). Only significant changes after Bonferroni correction for multiple comparisons are shown. **D**, Proportions of tumor, stromal and immune components in TME for left and right-sided tumors, for metastatic and primary tumors, for MSI high and MSI low tumors. Enrichment analysis for left-sided compared to right-sided tumor cells **(E),** for metastatic compared to non-metastatic tumor cells **(F),** for MSI high and MSI low tumors (**G**); red color indicates fold-change increase, blue color indicate the decrease of mean change enrichment score.

Post-treatment cohort included 24 biopsies (1 biopsy per patient) collected on average 100.2 (SD=105.3) days after the last day of therapy (Figure 1A). 15 patients, which demonstrated partial or complete tumor response to chemotherapy, and 3 patients with stable disease were categorized in this study as responders; and 6 patients had a progressive disease at the time of the biopsy (progressors) (Table 1). Majority of the tumors from this cohort were of high grades (stage III-IV, 22/24) and had a *KRAS* mutation (14/24). All four CRC molecular subtypes (CMS) were represented in the cohort (CMS1 – 6/24, CMS2 – 6/24, CMS3 – 3/24, CMS4 – 9/24) (Supplementary Fig. S1B,C). Most of the biopsies were taken from metastases (22/24). All those patients were treated with canonical chemotherapy FOLFIRI/FOLFOX and some of them received anti-VEGF or anti-EGFR agent (avastin, bevacizumab or cetuximab, respectively).

**Table 1.** Clinical characteristics of post-treatment cohort.

| Clinical characteristics | Values |
| --- | --- |
| Age, median (range) | 59.0 (39-78) years |
| Sex, n (%) |  |
| Male | 13 (54.2%) |
| Female | 11 (45.8%) |
| Primary tumor location, n (%) |  |
| Sigmoid | 6 (25%) |
| Right colon | 2 (8.3%) |
| Left colon | 8 (33.3%) |
| Rectum | 3 (12.5%) |
| Caecum | 1 (4.2%) |
| Rectum (high part) | 1 (4.2%) |
| Rectum (mean/lower part) | 2 (8.3%) |
| Rectosigmoid | 1 (4.2%) |
| Tumor type, n (%) |  |
| Primary | 2 (8.3%) |
| Metastasis | 22 (91.7%) |
| Response status, (RECIST), n (%) |  |
| PR | 14 (58.3%) |
| CR | 1 (4.2) |
| SD | 3 (12.5%) |
| PD | 6 (25%) |
| Treatment<br>type, (RECIST), *n* (%) |  |
| FOLFIRI | 4 (16.7%) |
| FOLFIRI + antiVEGF | 12 (50%) |
| FOLFIRI + antiEGFR | 4 (16.7%) |
| FOLFOX + antiVEGF | 4 (16.7%) |

We performed the scRNA-seq for those samples and obtained data for 463,630 single cells. Leiden clustering-algorithm was used to obtain cell type clusters (including tumor cells, different types of immune cells, and stromal cells), which were visualized on UMAP for each sub-cohort. Chromosomal instability scores were used to identify tumor cells and distinguish them from normal epithelial luminal cells (Figure 1B, Supplementary Fig. S2). With this approach, genomic instability was detected in all studied tumors. For a subset of tumors copy number profiles were validated with bulk WES (Supplementary Fig. S3). The average tumor purity in the cohort was 21.7% (SD=15.7%). Tumor cell populations were further analyzed using meta-pathway signatures from Gavish A. et al.[21], which revealed numerous differences in several pathways between studied subgroups of CRC. While characterizing the subgroups we considered tumor stage, *KRAS* mutational status and CMS group as covariates.

As a validation scRNA-seq cohort for untreated CRC we used single-cell treatment-naïve cohort of CRC adenocarcinomas from Pelka et al. (GSE178341), which included 26 MSS and 28 MSI, 13 left-sided and 41 right-sided tumors [21, 23].

### Left-sided CRC showed a greater propensity for a mesenchymal phenotype and low immunogenicity compared to right-sided CRC

In order to be able to dissect the impact of tumor location in the post-treatment cohort we first assessed the differences between left-sided and right-sided treatment-naïve primary MSS CRC. scRNA-seq resulted in 212,828 and 62,696 cells in left-sided and right-sided tumors. The side of the tumor location was not correlated with the tumor purity (23.2% (SD=15.9%) vs 15.6% (SD=12.3%)) nor with the overall fractions of stomal or immune components of the TME (Figure 1C-left). However within the immune cell compartment we observed significant increase in the number of B cells (p < 0.01) in right-sided compared to left-sided tumors (Figure 1D,E-left). Concordantly, in the scRNA-seq validation cohort we also revealed higher amount of plasma cells in right-sided compared to left-sided tumors (p < 0.01) (Supplementary Fig. S4).

Analysis of tumor cells using meta-pathway signatures[21] revealed significant differences in several pathways between right-sided versus left-sided CRC. We observed enrichment of Interferon/MHC-II signature (MP17) associated with the antigen-presentation in right-sided tumors (Figure 1G-left), which is in line with previously described higher antigenic load of right-sided tumors[15]. PDAC-classical signature (MP30) characteristic of mucinous phenotype in various cancers was enriched in right-sided CRC. This is consistent with data that the mucinous subtype of CRC is described to arise more often from right-sided tumors[24]. Left-sided tumors were characterized by increased respiration (MP21), cell cycle HMG-rich (MP3) and MYC pathway (MP20). In the scRNA-seq validation cohort, we confirmed increase of respiration (MP21), cell cycle HMG-rich (MP3) and MYC pathway (MP20) in left-sided tumors and PDAC-classical signature (MP30) and Interferon/MHC-II signature (MP17) in right-sided tumors as well as increased number of plasma cells in right-sided tumors compared to left-sided tumors (Supplementary Fig. S4).

#### Reduction in the number of plasma and B cells in treatment-naïve metastatic tumors versus primary tumors

In order to be able to separate the effects caused by treatments from the effects associated with metastasis in the post-treatment cohort we decided to investigate differences between metastases and primary tumors in treatment-naïve CRC. There was a decreased number of T helper 1 cells (p < 0.01), and plasma cells (p < 0,01) in metastatic cancers (Figure 1D,E-middle, Supplementary Fig. S5). These data accentuate the role of those cell types in suppression of metastasis, which was previously suggested for liver metastasis in CRC[25].

Metastatic cancer cells demonstrated enrichment of respiration (MP21), HMG-rich cell cycle (MP3) and MYC pathway (MP20), as well as proteasomal degradation (MP8) and EMT-related pathways (Figure 2F-middle), which was similar to observed differences between right-sided and left-sided tumors. Non-metastatic cancer cells were characterized by enhanced signatures related to the increased secretion of cytokines and chemokines (MP22, MP23) (Figure 2G-middle). Since EMT is known to be linked to metastasis in CRC, activated EMT-related pathways in left-sided colon might suggest increased metastatic propensity[26]. Indeed, in our extended cohort of 114 patients with detailed clinical annotations we revealed higher frequency of metastases and higher number of sites of metastasis in left-sided compared to right-sided tumors at the moment of diagnosis (p < 0.01, p < 0.05) (Supplementary Table S2). This is consistent with observations that left-sided colorectal cancer has a higher risk of liver metastasis[27, 28]. In addition, we found enrichment of CMS2 and CMS4 subtypes in left-sided tumors compared to right-sided tumors (p < 0.05) (Supplementary Fig. S1), this is also consistent with the data on the more frequent metastasis of the CMS4 subtype[29].

**Figure 2.**
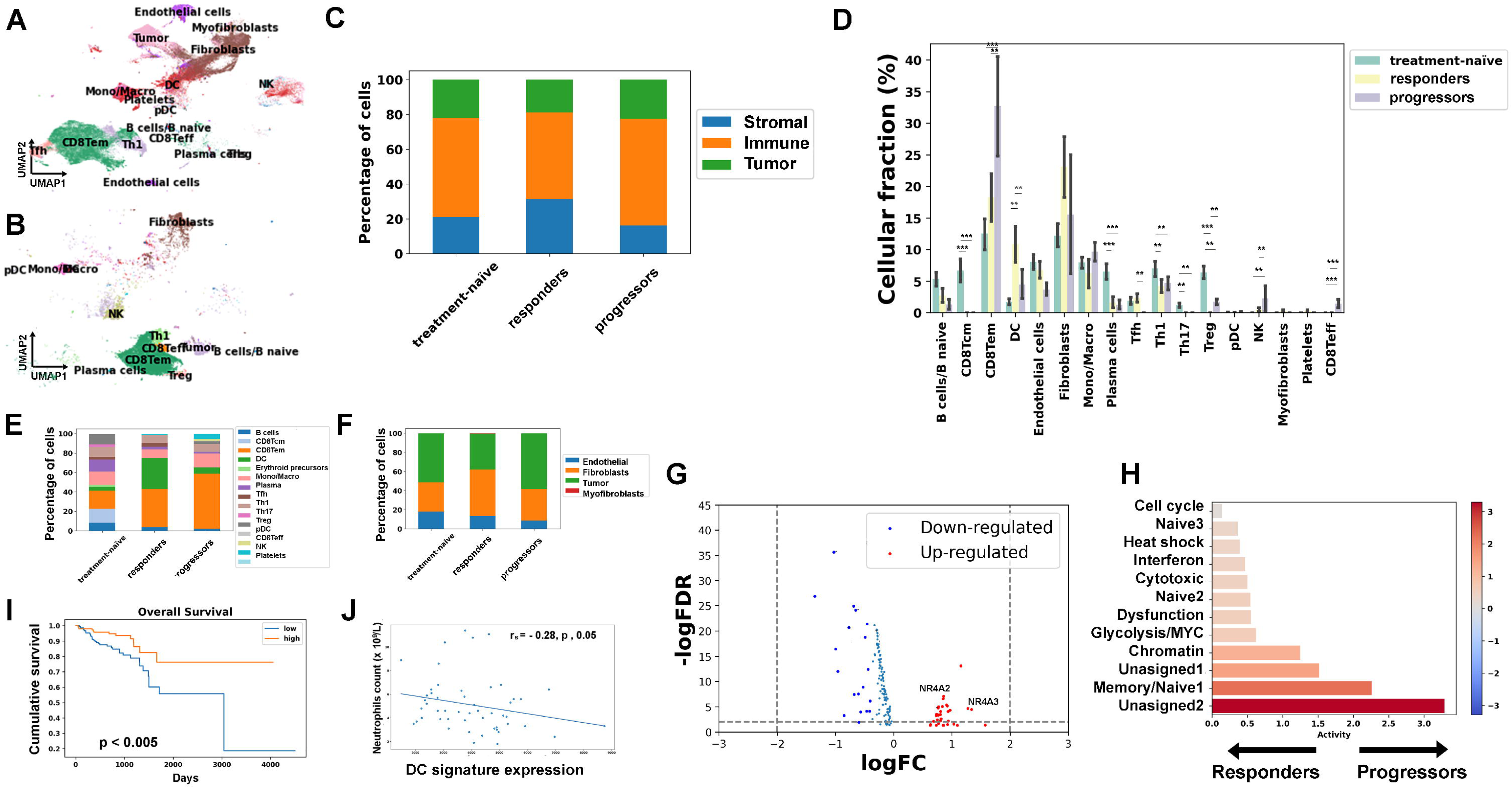
Characteristics of CRC after chemotherapy. Uniform manifold approximation and projection (UMAP) embedding of 146,746 cells (n = 18) from responders **(A)** and 31,276 (n = 4) cells from progressors **(B)**. **C**, Proportions of tumor, immune and stromal cells in treatment-naïve, responders and progressors. **D**, Barplots for differences in cell types proportions between treatment-naïve, responders and progressors. Whiskers indicate 75 confidence intervals and the asterisk represents p-values for the Mann-Whitney U test (* – p < 0.5, ** – p < 0.01, *** – p < 0.001). Only significant changes after Bonferroni correction for multiple comparisons are shown. Percentage of immune cells **(E)** and stromal cells **(F)** for treatment-naïve, responders and progressors. **G**, Differentially expressed genes in CD8+ effector memory T cells of progressors versus responders. **H**, Enrichment analysis for CD8+ T cells in progressors compared to responders. Kaplan-Meier curves for DC signatures in responders **(I)**. **J**, Correlation between DC signature and neutrophils counts in META-PRISM COADREAD cohort.

### MSI CRCs were more immunogenic and contained more Treg cells compared to MSS

We decided to investigate the extent of heterogeneity between MSS and MSI CRC. Overall, in the untreated cohort we identified of 35,297 tumor cells, including 30,387 from MSS tumors, and 4,910 from MSI tumors. In line with the expectations MSI tumors demonstrated lower tumor purity with a mean of 10.6% (SD=10.9%) vs 14.7 (SD=12%; Mann–Whitney U test, p < 0.05) for MSS tumors as well as an increased representation of Immune component[30, 31] (Figure 1C-right). This underpins the value of single-cell ­­analysis allowing considering pure cell type populations for the analysis. MSI tumors were different from MSS tumors by some aspects of tumor cell biology and by TME composition. At the level of TME MSI tumors were characterized by increased levels of Tregs, Th17 (p < 0.001, p < 0.01, respectively; Figure 1D,E-right), while MSS tumors had increased levels of Tfh, Th1 and pDC (Mann–Whitney U test, p < 0.001 for each) (Figure 1D,E-right). Concordantly, previous study revealed that MSI tumors were characterized by decreased numbers of CD4 T helper cells and significantly increased numbers of γδ T cells, a cell type, which is similarly to Tregs, has immune regulatory functions[32]. Stromal component of TME on MSI was characterized by increased infiltration of fibroblasts (Figure 1F-right), which formed two separated clusters (Figure 1B) in both, our untreated cohort and scRNA-seq validation cohort (Supplementary Fig. S4F).

Comparison of MSI and MSS tumors using meta-pathway signatures[21] revealed the strongest up-regulation of signature associated with Interferon/MHC-II in MSI tumor cells[33] (Figure 1G-right). PDAC-classical signatures (MP30) associated with mucinous phenotype was also enriched in MSI CRC. Analysis of the scRNA-seq validation cohort confirmed increased activity of both signatures in MSI CRC (Supplementary Fig. S4).

Considering the observed differences between MSS and MSI tumors in the untreated cohort we decided to include in the analyses of post-treatment CRC, which are represented by only MSS tumors in our cohort only MSS untreated tumors.

### Characteristics of CRC treated with chemotherapy

In order to reveal distinct features associated with response and progression after chemotherapy in CRC we compared those two sub-cohorts and used treatment-naïve MSS tumors as a baseline where it was appropriate. From scRNA-seq of post-treatment CRC we obtained 146,746 single cells from responders and 31,276 cells from progressors (Figure 2A,B). The mean tumor purities (TP) were 22.8 (SD=18.2) for responders and 21.2 (SD=11.5) for progressors. Analyses of the CMS compositions revealed increase of CMS4 in post-treatment tumors, which were predominantly metastatic versus mainly primary baseline MSS tumors (p < 0.05). At the same time, no significant differences in CMS compositions were observed between responders and progressors (Supplementary Fig. S1).

The decrease of plasma cells (p < 0.001), as well as of Th1 and Th17 cells (both p < 0.01), was observed in both subgroups of post-treatment CRC (Figure 2D,E). In responders there was a decrease of Treg cells (both p < 0.01) and increase of dendritic cells (DC), as compared to both treatment-naïve and progressors (both p < 0.01) (Figure 2D,E). At the same time, the higher abundance of immune cells in the TME was observed in progressors as compared to responders and treatment-naïve tumors (Figure 2C-E, Supplementary Fig. S6) with the particular enrichment of CD8 T effector, CD8 T effector memory and NK cells infiltration (p < 0.01, p < 0.001, p < 0.01, respectively). Contrary to expectations, in progressors the most prevalent fraction of TME cells was represented by CD8 T, CD8 T effector and effector memory T cells, which are usually associated with cytotoxicity[34] (Figure 2D,E). Interestingly, analysis of the CD8 T cells in progressors versus responders by Wilcoxon test with and without sample size correction revealed markers of the exhaustion phenotype - *NR4A2*, *NR4A3* among top 10 differentially expressed genes (DEG) [35] (Figure 2G). In line with this, analysis of meta-signatures of T cells (Gavish et al.) has also shown enrichment of signature associated with dysfunction in CD8 T cells of progressors compared with responders (Figure 2H).

### Antitumoral role of dendritic cells in CRC after chemotherapy

As DC were specifically enriched in responders, we decided to investigate their gene expression signature. The signature included 10 DEGs whose expression was specific to DC and that were overexpressed in responders vs. progressors. Survival analysis on 300 untreated tumors from COADREAD TCGA dataset (of colon and rectal cancer patients) revealed that high activity of DC-specific signature (*HLA-DPB1, FTL, FTH1, RPL10, TPT1, TMSB4X, RPL13, TMSB10, EEF1A1, RPS12*) was associated with better overall survival OS (HR = 0.56 CI = 95%, *p* < 0.005) (Figure 2I). This may indicate that the increased number of DCs might predict sensitivity to chemotherapy. Furthermore, we observed negative correlation between DC signature and blood neutrophils (r_s_ = -0.28, p < 0.05), a known marker of poor prognosis in a cohort of 51 metastatic refractory to conventional chemotherapy META-PRISM COADREAD tumors[36] (Figure 2J).

### Pericyte-like fibroblast subtype was associated with CRC aggressiveness

Fibroblasts in our single-cell analysis were grouped within two distinct clusters on UMAP projection for treatment-naïve and post-treatment sub-cohorts, which is in line with previous report on CRC[10] (Figure 3A). We decided to characterize those clusters, which we called FC1 and FC2, and to test their potential association with response to chemotherapy.

**Figure 3.**
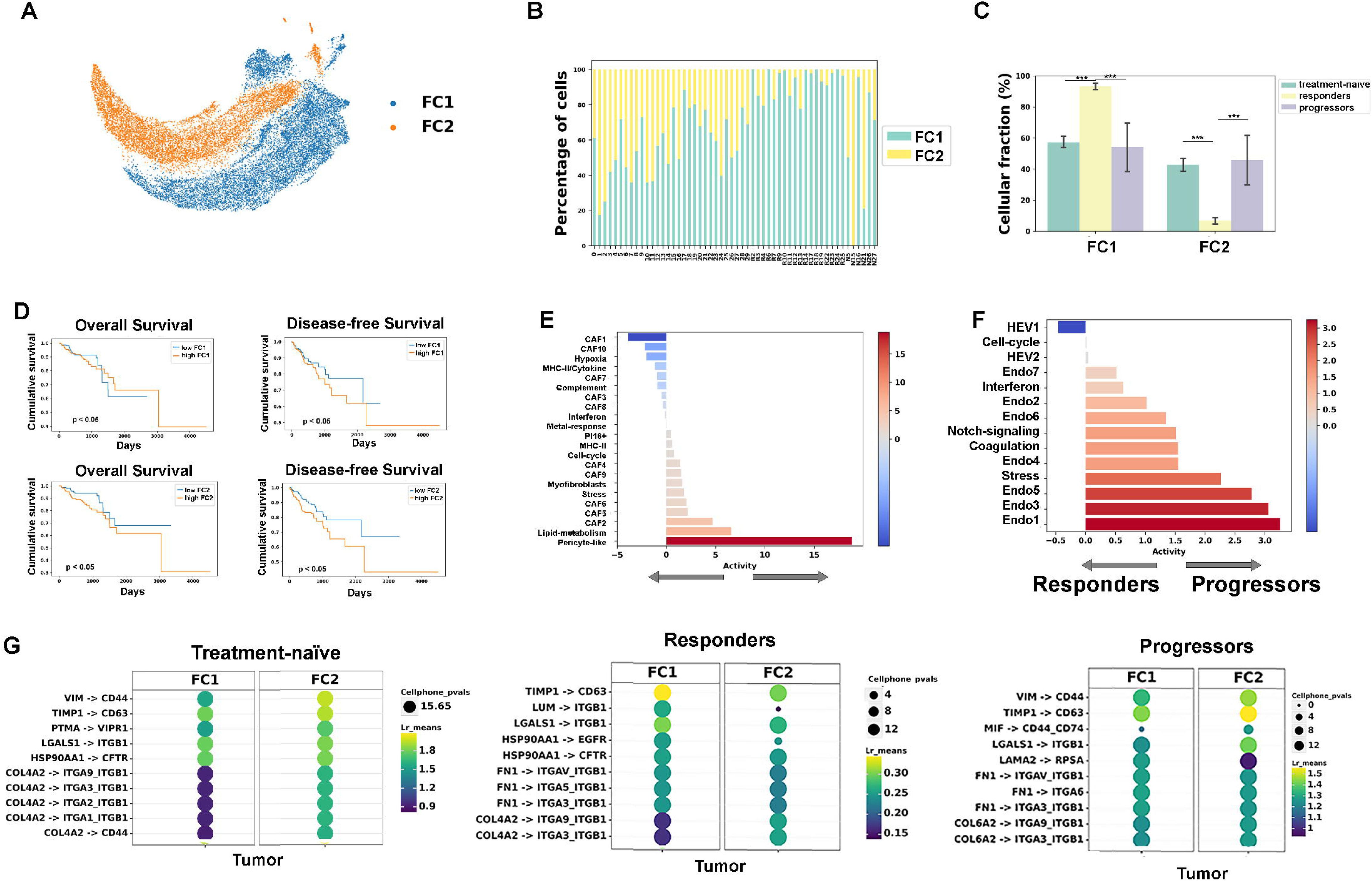
Proportions of fibroblast subtypes in treatment-naïve, response and progressors. Two separated clusters of fibroblasts in treatment-naïve (**A**). Changes in two subtypes of fibroblasts in treatment naïve, responders and progressors (**B**). Differences of FC1 and FC2 proportions for each sample (**C**). Kaplan-Meier curves for FC1 and FC2 gene signatures in TCGA COADREAD cohort (**D**). Enrichment analysis for FC1 and FC2 fibroblasts (**E**) and endothelial cells in responders compared to progressors (**F**). Ligand-receptor interactions between FC1/FC2 fibroblasts (source) and tumor cells (target) in treatment-naïve, responders and progressors (top 10 interactions) (**G**).

Specific gene signatures for FC1 (n=12 genes) and FC2 (n=13 genes), which included DEGs uniquely expressed in fibroblasts, were determined on untreated cohort (Figure 3A, Supplementary Table S3). These signatures were also delineating cluster separation in post-treatment cohort (Figure 3B). We compared those clusters with the two fibroblast subtypes (CAF-A and CAF-B) identified in Li et al, and observed a correspondence of FC1 to CAF-A, and of FC2 to CAF-B [10]. FC2 cluster in responders was significantly under-represented as compared to progressors and treatment-naïve tumors (Mann–Whitney U test, p < 0.001 for both) (Figure 3B,C). We then hypothesized that FC2 fibroblasts might be associated with aggressive phenotype in CRC. To test that we grouped samples from COADREAD TCGA cohort based on activity of these fibroblast signatures. The signatures of two fibroblast subtypes had different effects on OS and progression-free survival (PFS) in this cohort. FC2 signature was associated with poor OS and PFS (1.22 and 1.19, p < 0.05 for both), while FC1 signature was not predicting OS or PFS (Figure 3D). Strikingly, meta-pathway enrichment analysis for fibroblasts revealed that FC2 was distinguished from FC1 by strong activity of pericyte-like signature (Figure 3E). Moreover, meta-pathway enrichment analysis for endothelial cells revealed that in responders there was an increased activity of high endothelial venules (HEV) pathway compared to progressors, while progressors had enriched signatures associated with endothelium (Figure 3F), which are known to be surrounded by pericytes[37].

In order to investigate interaction between different cell types in TME and tumor cells we performed ligand-receptor interaction analysis independently for the three groups: treatment-naïve, responders and progressors. Among the 30 strongest interactions across groups, the interaction between FC2 and tumor cells via VIM-CD44 was observed only in progressors and treatment-naïve tumors (Figure 3G). This may indicate that CD44+ cancer stem-like cells recruit FC2 mesenchymal fibroblasts in treatment-naïve tumors and progressors, but not in responders. This is in line with our previous finding that responders are characterized by a decrease of FC2 fibroblast subtype, which is associated with poor survival. The poor effect on survival of the FC2 subtype may be due to its association with its pericyte function and potentially vasculature.

### Chemoprotective function of anti-apoptotic genes, expressed in stem-like tumor cells in progressive disease after chemotherapy

In order to understand how tumor cells react to chemotherapy, we focused the following analysis only on a tumor cells population (Supplementary Fig. S2). We first selected tumor cells from treatment-naïve CRC, responders and progressors (35,297, 27,571 and 9,269 cells respectively) (Figure 4A). UMAP projection without integration resulted in separate clusters for each tumor. As we expected substantial differences in transcription profiles between tumor samples were observed; this might be due to different profiles of Somatic Copy Number Alterations and other genomic and epigenetic features, so we did not apply batch correction at re-clusterization (Figure 4A). Among the top 10 most significant overexpressed DEGs in progressors compared to responders there was an anti-apoptotic gene *MTRNR2L1* (p < 0.001) (Figure 4B-D). Among top 30 genes, there was also *CDX1*, known by its protection function for cancer stem cells in colorectal cancer (p < 0.001) (Figure 4B-D)[7]. We found that there was a significant and strong correlation between these two protective genes with markers of stemness *CD24* and *EPCAM*, specifically in progressors (r_s_ =0.43−0.46), while it was weaker in responders (r_s_ =0.15−0.25) and treatment-naïve tumors (r_s_ =0.066−0.23) (Figure 4C,D). In order to test for potential batch effect from different samples, we repeated the analysis with batch correction at re-clusterization, and the results for all the three identified genes remained significant. In a META-PRISM COADREAD cohort we showed the positive correlation between mean expression of *CDX1*, *MTRNR2L1* and *CD24* signature and Gustave Roussy Immune Score (GRIm), which is known by association with poor survival and inflammation in CRC[36, 38] (r_s_ = 0.38, p < 0.001) (Figure 4E).

**Figure 4.**
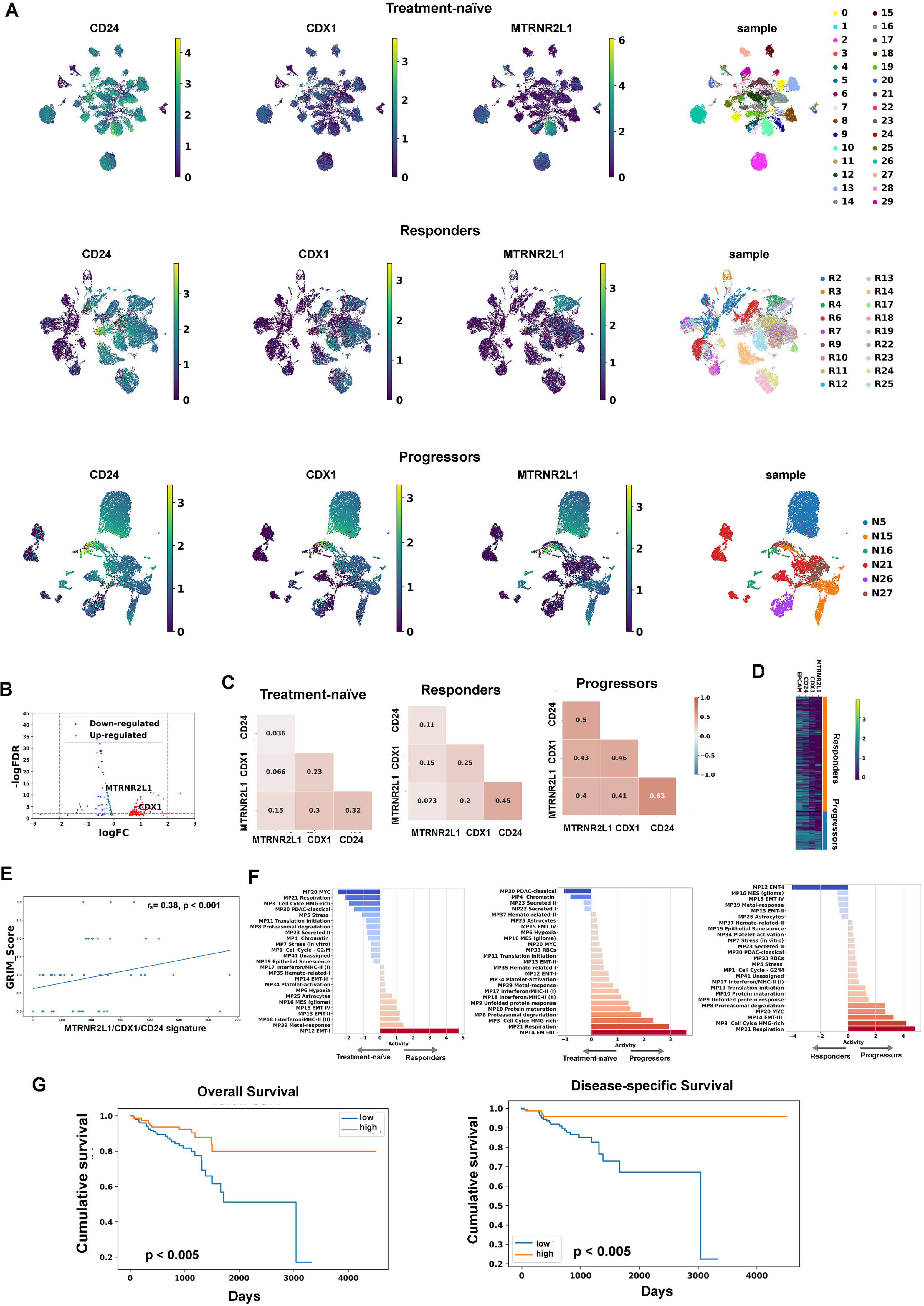
Differences in expression of CD24, CDX1 and MTRMR2L1 in cancer cells before and after chemotherapy in responders and progressors. **A**, UMAP plots after leiden re-clustering of 33,911 cancer cells from 30 treatment naïve samples, 27,571 cancer cells from 18 responders and 9,269 tumor cells from 6 progressors. **B**, Differentially expression analysis for progressors and response cancer cells. Volcano plot for DEGs in progressors compared to responders. Spearman correlation of *CD24*, *CDX1* and *MTRMR2L1* in treatment-naïve, responders and progressors **(C)**. The heatmap for all responder and progressors cells shows the expression of *CD24*, *CDX1*, and *MTRNR2L1* in each cell **(D)**. Spearman’s correlation between MTRNR2L1, CDX1 and CD24 signature and GRIM score in a META-PRISM COADREAD cohort **(E**). Enrichment analysis for cancer cells, mean change. Red bars shows mean change for increased pathways, blue bars shows mean change for decreased pathways (p < 0.05) **(F)**. Kaplan-Meier plots for signature associated with cancer cells in responders **(G)**.

Meta-pathway enrichment analysis revealed up-regulation of signatures associated with respiration, HMG-rich cell cycle, MYC pathway and EMT-III, which includes both mesenchymal and epithelial markers, consistent with a hybrid cellular state in progressors, as compared to treatment-naïve tumors and responders (Figure 4F). There was also an up-regulation of both signatures, associated with interferon and MHC-II immune response (MP17, MP18) in progressors compared to responders, but not to treatment-naïve tumors (Figure 4F). In responders there was a significant upregulation of EMT-I signature, representing full mesenchymal phenotype[21], while MYC and G2M signatures were down-regulated (Figure 4F).

We further identified a set of genes characterizing tumor cells in therapy response. We selected strongest tumor cell-specific DEGs that were not expressed in cells of TME (*KRT18*, *PTPRK*, *KRT19*, *PHGR1*, *LLGL2*, *LMO7*, *CTNND1*, *AGAP1*, *MAP7*, *LINC00511*). This gene signature had a prognostic value in COADREAD TCGA cohort and was associated with significantly better OS and PFS (HR = 0.8, CI = 95%, p < 0.005 and HR = 0.52, CI = 95%, p < 0.005, respectively) (Figure 4G).

### Chemo-resistant apoptotic tumor cells express chemokines

We hypothesized that in post-treatment tumors responders might differ from progressive disease by the abundance of apoptotic cells and mechanism of apoptosis. In order to study this we performed an enrichment analysis for the HALLMARK_APOPTOSIS (M5902) signature from MSigDb (v2023.2) and calculated the average score for each cell[39]. We then found a list of genes that correlated with the score of apoptosis. Remarkably, chemokines *CXCL1*, *CXCL8*, *CXCL2*, and *CXCL3* were the top 4 genes correlated with apoptosis signature in progressors, but not in responders or treatment-naïve tumors (Figure 5A,B). They were also correlated between each other, particularly in progressors (p < 0.001) (Figure 5C-E). This signature was absent among the top 30 genes associated with apoptosis in responders. Among other genes correlated with apoptosis signature in progressors there were NF-κB pathway genes: *RELB* and *NFKBIA* (Supplementary Table S4). On the contrary *CDKN1A*, associated with cell cycle arrest and DNA damage response, had the highest correlation with apoptotic signature in responders and absent among top 30 genes in progressors and treatment-naïve tumors[40] (Supplementary Table S4, Figure 5A). This was in line with meta-pathway analysis, which showed downregulation of G2M pathway in responders. There was also a progressor-specific negative correlation of *CXCL8* with *CD24*, *MTRNR2L1* and *CDX1* (r_s_ = -(0.04 – 0.17), p < 0.05) (Figure 5E), which is in line with our hypothesis that stem-like cells expressing protective markers less frequently undergo apoptosis then other tumoral cells. Thus, in progressors apoptosis appears to be more associated with inflammation and chemokines expression, while in responders apoptosis pathway seems to be more associated with mitotic stress.

**Figure 5.**
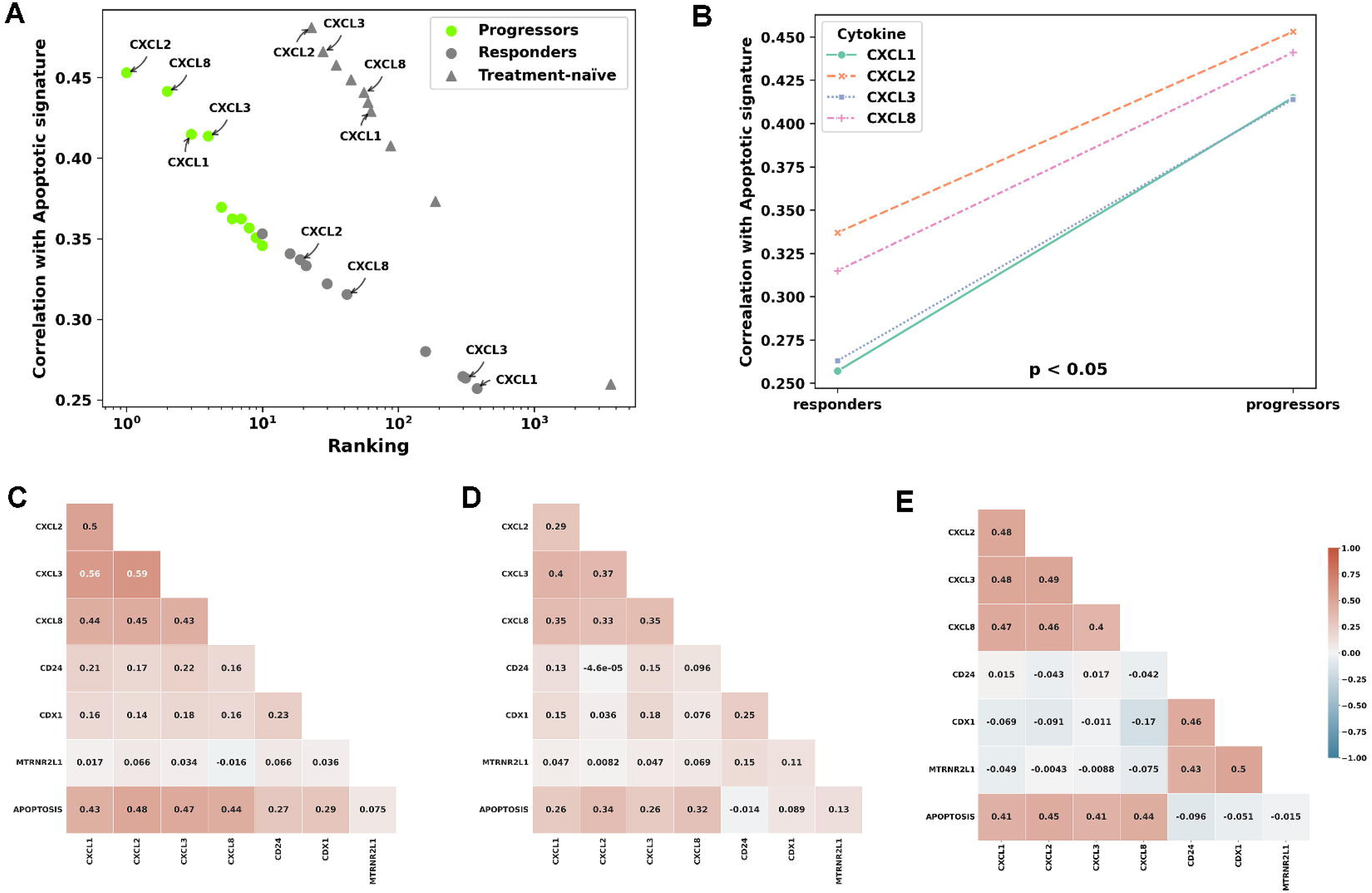
The major changes in apoptotic chemo-resistant cells are related to the expression of chemokines. Top 10 genes positively correlated with the HALLMARK_APOPTOSIS signature in progressors displayed for treatment naïve (gray triangle) and responders (gray circles), p < 0.001 for the Spearman rank correlation coefficient (**A**). Differences in correlation coefficients with the apoptotic signature for four CXCL cytokines between responders and progressors, p-value for the Mann-Whitney test is shown (**B**). Heatmap shows changes in Spearman’s correlation coefficient from -1 (blue) to 1 (red) for the genes CXCL8/1/2/3, MTRNR2L1, CDX1 and CD24 with the HALLMARK_APOPTOSIS signature in treatment-naïve (**C**), responders (**D**), and progressors (**E)**.

## DISCUSSION

This is a study of a large scRNA-seq cohort of 56 CRC, which uncovers associations of tumoral cells and TME composition with response and progression after treatment with conventional chemotherapy. Analyses of treatment-naïve CRC cohort is in line and provides additional precision to the expected molecular characteristics of tumor cells and peculiarities of TME associated with clinical patterns. In particular, we found that right-sided CRC exhibited high immunogenicity, which followed from TME composition: increased counts of B cells; and from tumor cells: active Interferon/MHC-II signature compared to left-sided CRC. We also observed that MSI tumors were associated with higher levels of immune cells in the TME despite high levels of Treg cells, which are thought to elicit immunosuppressive function. Though Tregs were characterized as marker of aggressive tumor in various cancer types[41], it was recently suggested that they might predict a good response to immunotherapy, which is actually the case of MSI CRC[42].

This work proves that scRNA-seq of post-treatment cancers helps to stratify cohorts by the markers of aggressiveness, prognosis and resistance to treatment. Specifically, we confirmed existence of two subtypes of fibroblasts in CRC, which were previously reported, and furthermore we were able to associate one of the two subtypes, FC2, with aggressive tumors. FC2 fibroblasts with enriched pericyte-like signature might be involved in vascularization and angiogenesis in progressors, which is congruent with our finding of enrichment of endothelium signatures in progressors. Remarkably, association of FC2 fibroblasts signature in independent bulk RNA-seq cohort from TCGA with poor survival suggests that it might be a perspective marker of aggressive CRC and of poor response to chemotherapy. On the other hand, signature of DC in responders was associated with good prognosis in CRC TCGA cohort, which included tumors subjected to treatment after the biopsy. Additionally, in META-PRISM cohort of advanced metastatic and refractory tumors we revealed negative association of DC signature with neutrophil levels, which is a known marker of poor prognosis and inflammation. In line with the fact that chemotherapy can enhance anti-tumor DC function[43], our results show important immune modulation in treatment response. Along with the signatures from the cells of the TME, signature derived from the tumoral cells, which was based on the DEGs specific to responders, was a strong prognostic factor in untreated CRC from TCGA.

This work also demonstrates potential for revealing specific mechanisms of resistance to chemotherapy in CRC. For example, progressors showed increased expression of the anti-apoptotic factors *MTRNR2L1* and *CDX1*, which are also associated with protection of tumor cells from cytotoxic effects of chemotherapy and with autophagy [7, 44]. These results together with co-expression of *MTRNR2L1* and *CDX1* with the stem cell marker and immune checkpoint *CD24*, propose a potential mechanism of chemotherapy resistance in CRC (Figure 6). Interestingly, inhibitors of *CD24* were shown to improve efficacy of chemotherapy in CRC[45]. Previously, it was described that stem cells can mediate chemoresistance through autophagy, which can help avoiding the cytotoxic effects of T cells [7, 46]. In other studies, on different cancer types it was also reported that stem cells are enriched in tumors with “hybrid” EMT phenotype represented by a mixture of epithelial and mesenchymal cells [47]. Our data concordantly demonstrated a prevalent “hybrid” EMT signature (EMT-III) in progressive CRC, which correlated there with expression of protective markers in stem-like cells.

**Figure 6.**
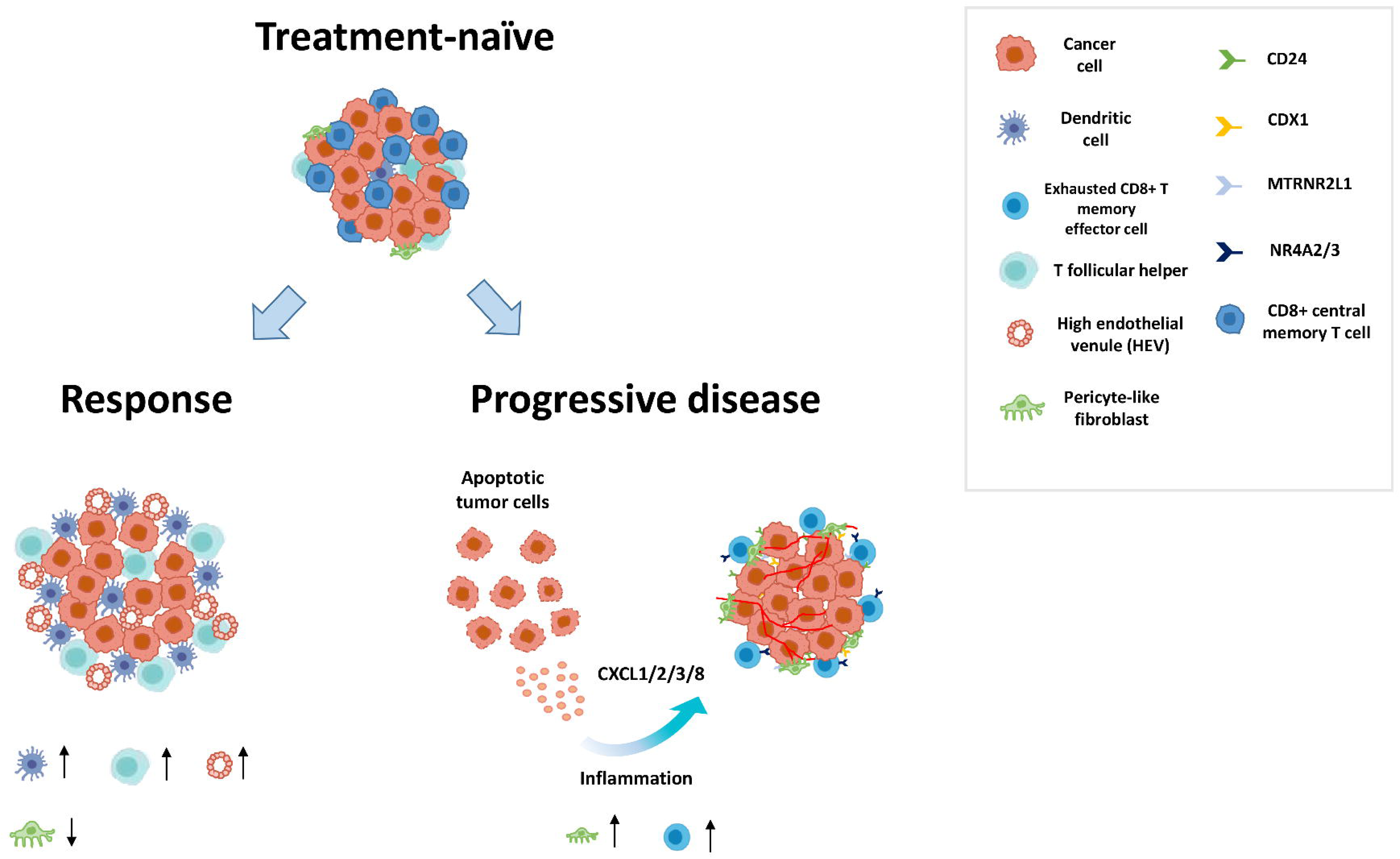
**Schematic illustration of main changes in TME and tumor cells in responders and patients with progressive disease after conventional chemotherapy.**

We leveraged scRNA-seq data to characterize state of different cell populations of the TME. For example, there was an elevated level of CD8 T-effector memory cells in progressors, which was not expected as this cell type usually elicits antitumoral function. High level of CD8 T-effector memory cells in progressors was in line with the finding that tumor cells in progressors showed activity of interferon and MHC-II signatures, which are associated with lymphocyte infiltration. However, analysis of transcriptional patterns in this cell population revealed an overexpression of markers of exhaustion, such as *NR4A2* and *NR4A3*, revealing that those cells are likely dysfunctional in progressors. This finding indicates a potentially pro-inflammatory TME phenotype in progressors[35]. This is in line with previous data that inflammation caused by chemotherapy can lead to decreased effectiveness of therapy[48]. Furthermore, anti-inflammatory agents such as aspirin have recently been shown to increase chemosensitivity in CRC, which also indicates a link between inflammatory responses and chemoresistance[49]. We have found that CXCL8 along with other CXCL chemokine family genes were among the top overexpressed genes in apoptotic cells in progressive disease after chemotherapy. This finding is in line with the previous study where it was shown on CRC cell lines treated with oxaliplatin, that CXCL8 chemokine is expressed by apoptotic tumor cells in CRC and is responsible for the pro-inflammatory TME[50].

While comparing post-treatment and baseline CRC is it important to discuss the impact of cohort heterogeneity that might have biased the results. We observed a decrease of plasma cells and Type 1 T helper cells (Th1) in post-treated tumors versus untreated tumors. This actually could be explained by the fact that in our cohort post-treated tumors were predominantly metastatic (Table 1), while treatment-naïve tumors were mainly primary. Indeed, similar decrease of these cell types was observed in metastatic tumors versus primary tumors if only untreated tumors are considered. At the same time, these were the only results of comparison of post-treatment and untreated tumors that could be explained by the impact of metastasis.

Overall, we identified markers of response to chemotherapy and of progressive disease at the level of TME and of tumor cells. Corresponding signatures were derived from tumor cells, CAFs and dendritic cells, and were shown to be associated with survival on large TCGA and META-PRISM cohorts. We also revealed that in cancer cells in progressive disease there was an overexpression of genes protecting cells from cytotoxicity. These genes were specifically expressed in cancer stem-like cells, suggesting existence of a protection mechanism for cancer stem cells against chemotherapy. Finally, we observed that apoptotic cells were associated with different signatures between responders and progressors. In responders, apoptotic cells demonstrated cell-cycle dysregulation, while in progressors they were characterized by activated signature of secretion of pro-tumoral chemokines (*CXCL1, CXCL2*, *CXCL3* and *CXCL8*). Those results suggest that inflammation is associated with progressive disease after chemotherapy.

In summary, our findings emphasize the impact of tumor cell signatures, fibroblast subtypes and dendritic cells on post-chemotherapy survival outcomes, highlighting their potential prognostic value, which has to be further validated in clinical studies. The elevated expression of resistance-associated markers in progressors and the immune landscape disparities between responders and progressors underscore the complexity of treatment responses in CRC, paving the way for personalized therapeutic strategies and a deeper understanding of immune modulation in cancer management. These intricate differences in immune cell compositions, fibroblast subtypes, molecular interactions, and immune exhaustion markers underpins the multifaceted nature of the TME in CRC treatment responses, providing valuable insights for tailored therapeutic strategies and patient management in colorectal cancer settings.

## METHODS

### Sample collection

The cohort includes 32 samples from untreated patients and 24 samples from patients who received chemotherapy. All patients were diagnosed with colorectal cancer between 2014 and 2022 at the Gustave Roussy Institute. Chemotherapy included FOLFOX or FOLFIRI as first line, in some cases in combination with anti-VEGF or anti-EGRF drugs (avastin, bevacizumab or cetuximab) as the second and third lines. The post-treatment cohort included 18 responders (partial response, stable disease and complete response) and 6 progressors (progressive disease). To validate the signatures found, we used a cohort from the META-PRISM dataset, which included patients with colorectal and rectal cancer treated with FOLFIRI or FOLFOX alone or in combination with anti-VEGF or anti-EGFR therapy[36]. All biopsies were taken after treatment.

### Single-cell data analysis

Single-cell data was analyzed according to a standard tutorial using the Scanpy package[51]. Cells were filtered by the percentage of mitochondrial (<15%) and ribosomal (<30%) genes. Cell doublets were also detected and removed using the Scrublet package (https://github.com/swolock/scrublet). The leiden algorithm was performed for cell clustering[52]. For Uniform Manifold Approximation and Projection (UMAP) graphs computing parameters of neighbors n_neighbors=10 and number of components n_pcs=40 were used. In the principal component, analysis ARPACK SVD solver was used. Annotation of cell types occurred using the Decoupler package[53], which included data from the PanglaoDB cell type marker genes database based on sing-cell expression data[54]. Signatures from TISCH2 were also used to determine T cell types[55]. Tumor cells were individually identified using the inferCNV package, based on the calculation of chromosomal instability in each cell. Data for GRCh38.p14 human genome from GENCODE were used to annotate the genomic location of genes. All immune cells were taken as reference. Data from different samples were integrated using the scvi-tools package, using batch correction with negative binomial distribution model for treatment-naïve, responders and progressors separately[56–58]. To visualize the results, the built-in Scanpy functions and the Matplotlib and Seaborn libraries for statistical data visualization were used. There were no significant differences in TME associated with tumor stage (TI/II vs TIII/IV, Mann–Whitney U test) and KRAS mutation (Supplementary Fig. S5). Consensus molecular subtypes (CMS) for CRC classification inferred from pseudo-bulk transcriptomes using CMScaller R package [29] (Supplementary Fig. S1). For convenience, we made summary tables of the changes found in treatment-naïve and post-chemotherapy cohorts (Supplementary Fig. S5, Supplementary Fig. S6).

### Defining gene signatures specific for cell types

After UMAP clustering and cell type annotation, we identified specific genes for each individual cluster corresponding to cell types. To do this, we performed a Wilcoxon test comparing each cell type with all other cell types combined. Then, to find genes unique to a particular cell type, we selected the top 25 differentially expressed genes and then selected only those of this genes that did not appear among the top 25 genes of any other cell type.

### Enrichment analysis

Over-representation analysis was performed to identify pathways altered in chemoresistant cells. We used the run_ora function of the decoupler package and signatures determined based on the integrated analysis of tumor single cell data by Gavish et al.[21]. Only labels with statistically significant changes were displayed on the graphs. To analyze the correlation between genes and signatures, the Spearman’s correlation coefficient was calculated using the Scipy package[59]. The CMScaller R package was used for consensus molecular subtyping or CRC cohorts[60].

### Differential gene expression analysis

The Wilcoxon test was used to analyze differential expression in progressors versus responder cancer cells. At the same time, down-sample simulations were used to take into account the heterogeneous number of cells in each sample. Down sampling for tumor cells was performed 5 times to confirm the reproducibility of the found differential expression. The Bonferroni test was used to take into account the correction for multiple losses. Changes with p-value less than 0.05 were considered statistically significant.

### Survival analysis

For survival analysis, TCGA COADREAD data from patients with colon and rectal cancer were downloaded. Clinical survival data and normalized FTPM gene expression RNA-Seq data for 728 patient were downloaded using the XENA browser[61, 62]. A cohort of patients who received primary therapy but did not receive radiotherapy was selected (n = 300). The Lifelines package was used to calculate Kaplan-Meier curves and hazard ratios (HR) (https://doi.org/10.5281/zenodo.1252342). The logrank test was used to compare survival curves. Changes with p-value less than 0.05 were considered statistically significant.

## Declarations

### Ethical Approval

All patients samples used in this study were stored in a tumor bank at the Gustave Roussy Cancer Campus. Written inform consent for the NGS was obtained from participants, as part of the following studies: MATCH-R (NCT02517892), INCB 54828-202 (NCT02924376), STING (NCT04932525), BoB (NCT03767075) or STARTRK-2 (NCT02568267).

## Author’s Contributions

**G. A. Puzanov:** Data curation, formal analysis, validation, investigation, visualization, methodology, writing–original draft, writing–review and editing. **C. Astier:** Resources, data curation. **A. A. Yurchenko:** Methodology, data curation. G. Jules-Clement: Resources, data curation. **F.Andre:** Resources. **A. Marabelle:** Validation. **A. Hollebecque:** Data curation, resources, validation. **S.I. Nikolaev:** Conceptualization, validation, methodology, data curation, supervision, writing–original draft, writing–review and editing.

## Availability of data and materials

Expression matrices with scRNA-seq data for the studied cohort are available upon request. FASTQ source files are available on https://ega-archive.org. RNA-Seq and clinical data for COAD and READ cohorts are available at the website of the TCGA project: https://portal.gdc.cancer.gov/projects/TCGA-COAD; https://portal.gdc.cancer.gov/projects/TCGA-READ. Standardized data for validation cohort from META-PRISM project are available at: https://cbioportal.gustaveroussy.fr/study/summary?id=metaprism_2023.

