## Supplementary Fig. S1 for "Single-cell Transcriptome Profiling of Post-treatment and Treatment-naïve Colorectal Cancer: Insights into Putative Mechanisms of Chemoresistance"

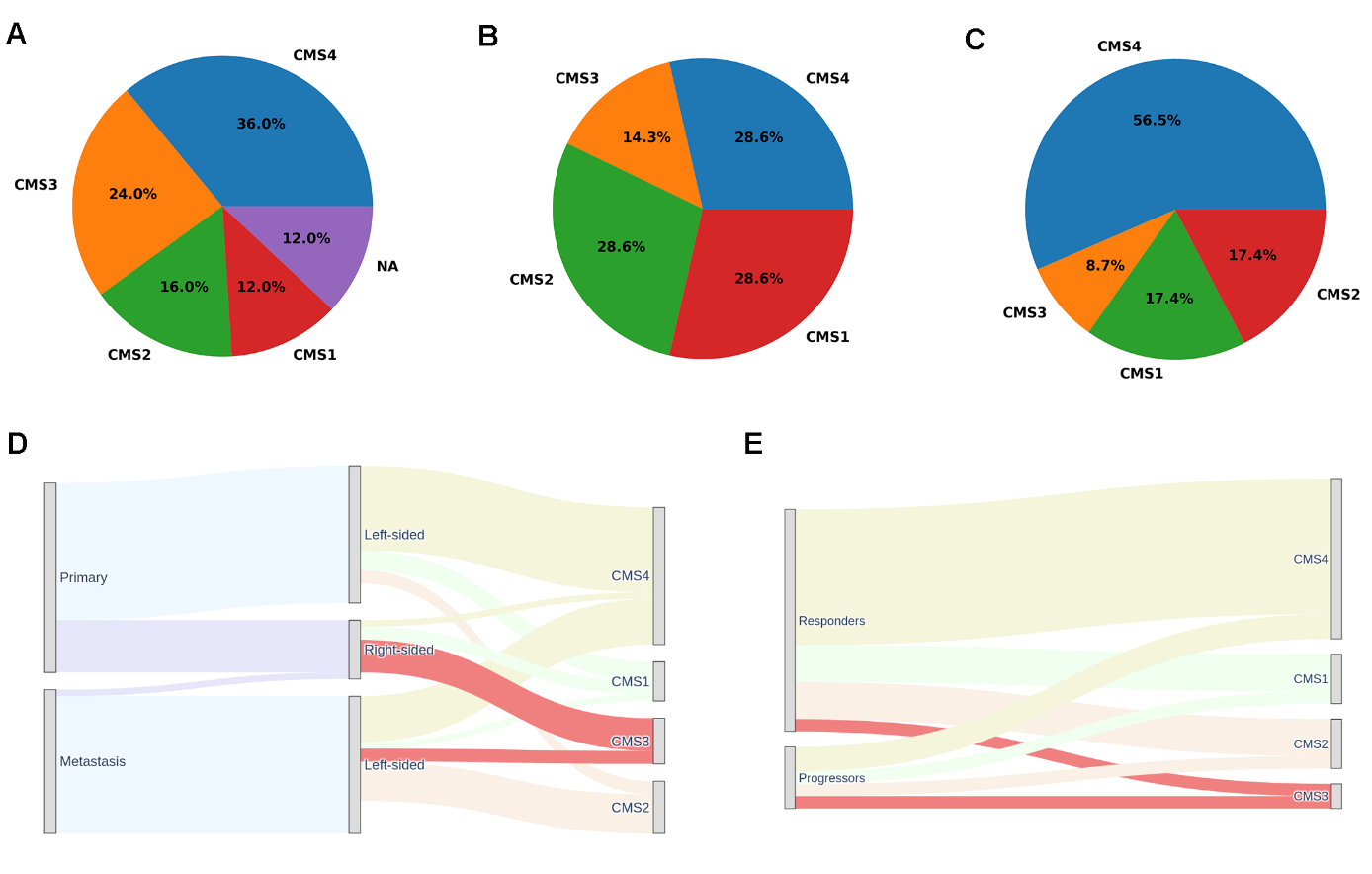

**Supplementary Figure S1. Predicted status of tumors using consensus molecular subtypes (CMS) for CRC classification.** Pie chart with proportions of CMS groups for untreated MSS (**A**), MSI (**B**) and post-treatment cohort (**C**). Sankey plots show the proportions of CMS subtypes in primary and metastatic tumors arising from left-sided and right-sided primary tumors (**D**) and in responding and progressive CRCs after chemotherapy (**E**).
