## Supplementary Fig. S2 for "Single-cell Transcriptome Profiling of Post-treatment and Treatment-naïve Colorectal Cancer: Insights into Putative Mechanisms of Chemoresistance"

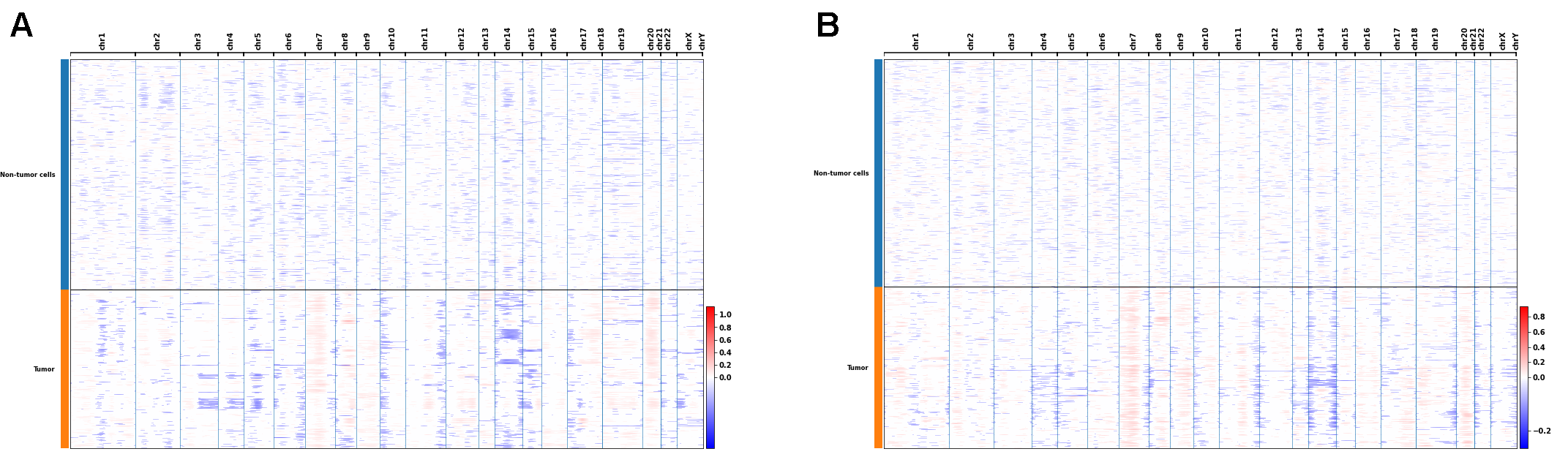

**Supplementary Figure S2. Identification of tumor cells by copy number variation.** Heatmap with CNVs for tumor cells in each sample of treatment-naïve (**A**) and post-treatment cohort with hierarchical relationship between the samples (**B**). Red indicates gain and blue indicates loss of chromosomal region.
