## Supplementary Fig. S6 for "Single-cell Transcriptome Profiling of Post-treatment and Treatment-naïve Colorectal Cancer: Insights into Putative Mechanisms of Chemoresistance"

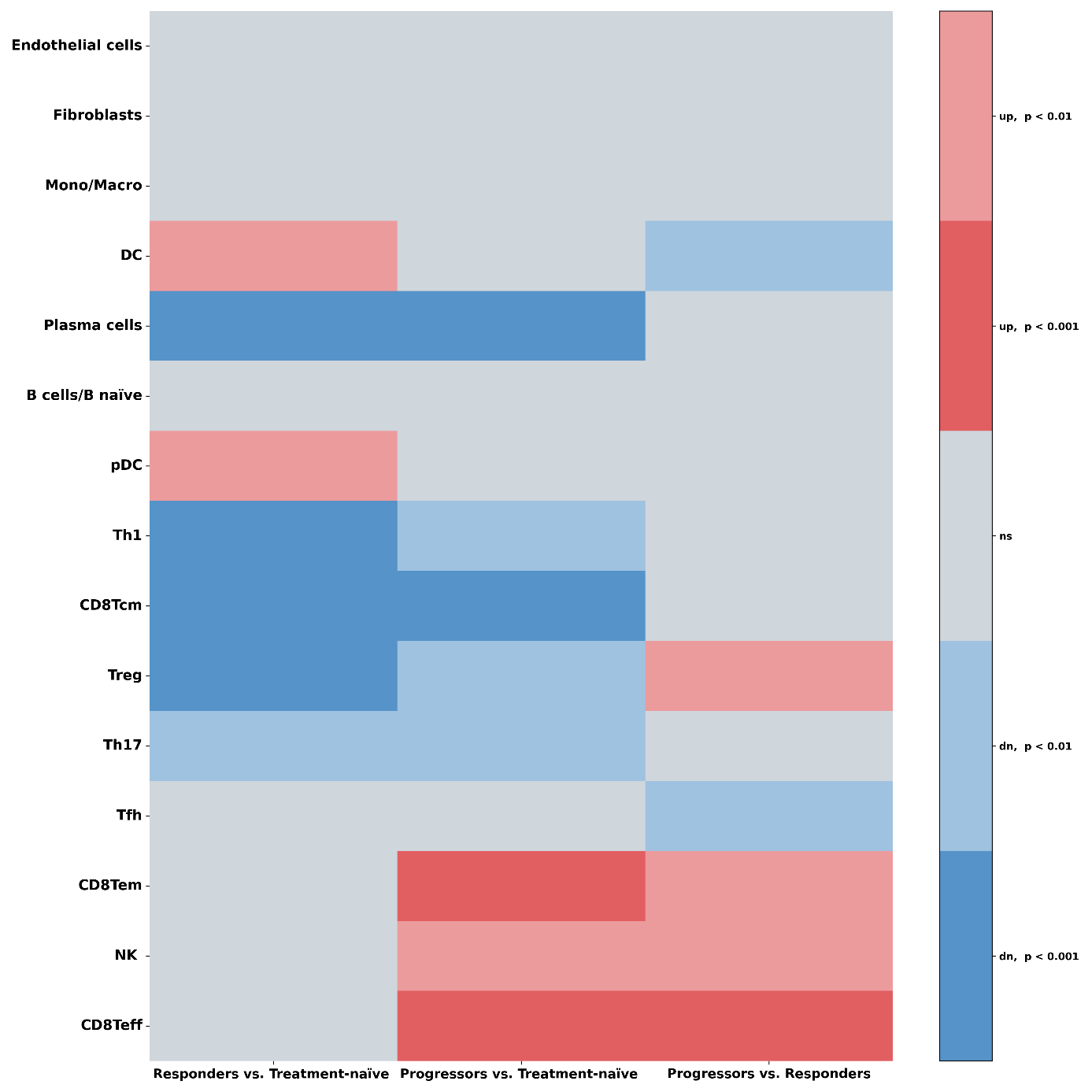

**Supplementary Figure S6. Changes in cell types proportions in TME of responders vs. progressors CRC arter chemotherapy.** Only significant changes after the Mann-Whitney test with Bonferroni correction for multiple comparisons are shown.
