## Supplementary Table. S1 for "Single-cell Transcriptome Profiling of Post-treatment and Treatment-naïve Colorectal Cancer: Insights into Putative Mechanisms of Chemoresistance"

Clinicopathologic features of treatment-naïve samples (MSS + MSI).

| **Clinical characteristics** | **Values** |
| --- | --- |
| Age, median (range) | 66.5 (23-90) years |
| Sex, *n* (%) | |
| Male | 14 (43.7%) |
| Female | 18 (56.3%) |
| Primary tumor location, *n* (%) |  |
| Sigmoid | 4 (12.5%) |
| Right colon | 6 (18.8%) |
| Transverse colon (left part) | 2 (6.3%) |
| Left colon | 8 (25.0%) |
| Rectum | 2 (6.3%) |
| Junction rectum-sigmoid | 2 (6.3%) |
| Caecum | 2 (6.3%) |
| Rectum (high part) | 4 (12.5%) |
| Rectum (lower part) | 1 (3.1%) |
| Rectosigmoid | 1 (3.1%) |
| Tumor type, *n* (%) |  |
| Primary | 27 (84.4%) |
| Metastasis | 5 (15.6%) |
