## Supplementary Table. S2 for "Single-cell Transcriptome Profiling of Post-treatment and Treatment-naïve Colorectal Cancer: Insights into Putative Mechanisms of Chemoresistance"

Clinical characteristics for extended cohort of CRC patients.

|  | **Left-sided** | **Right-sided** |
| --- | --- | --- |
| **Age** | 62.5 | 61.1 |
| **Female** | 40 | 14 |
| **Male** | 51 | 9 |
| **Metastasis status** | | |
| **Primary** | 50 | 17 |
| **Metastatic** | 41 | 6 |
| **Site of metastasis** | | |
| **Liver** | 39 | 4 |
| **Lung** | 13 | 2 |
| **Peritoneal carcinomatosis** | 10 | 3 |
| **Lymph node** | 10 | 1 |
| **Bone** | 2 | 0 |
| **Other** | 3 | 0 |
