## Supplementary Table. S3 for "Single-cell Transcriptome Profiling of Post-treatment and Treatment-naïve Colorectal Cancer: Insights into Putative Mechanisms of Chemoresistance"

Specific markers for FC1 and FC2 subtypes of fibroblasts.

| **gene** | **subtype** |
| --- | --- |
| COL6A3 | FC1 |
| GPC6 | FC1 |
| RARRES2 | FC1 |
| DCN | FC1 |
| BICC1 | FC1 |
| LUM | FC1 |
| C1S | FC1 |
| COL5A1 | FC1 |
| ANTXR1 | FC1 |
| COL12A1 | FC1 |
| VCAN | FC1 |
| CDH11 | FC1 |
| COL18A1 | FC2 |
| SOX5 | FC2 |
| EBF1 | FC2 |
| DLC1 | FC2 |
| RGS5 | FC2 |
| PDGFRB | FC2 |
| LHFPL6 | FC2 |
| CACNA1C | FC2 |
| FILIP1L | FC2 |
| CCDC102B | FC2 |
| NOTCH3 | FC2 |
| MGP | FC2 |
| RBMS3 | FC2 |
