## Supplementary Table. S4 for "Single-cell Transcriptome Profiling of Post-treatment and Treatment-naïve Colorectal Cancer: Insights into Putative Mechanisms of Chemoresistance"

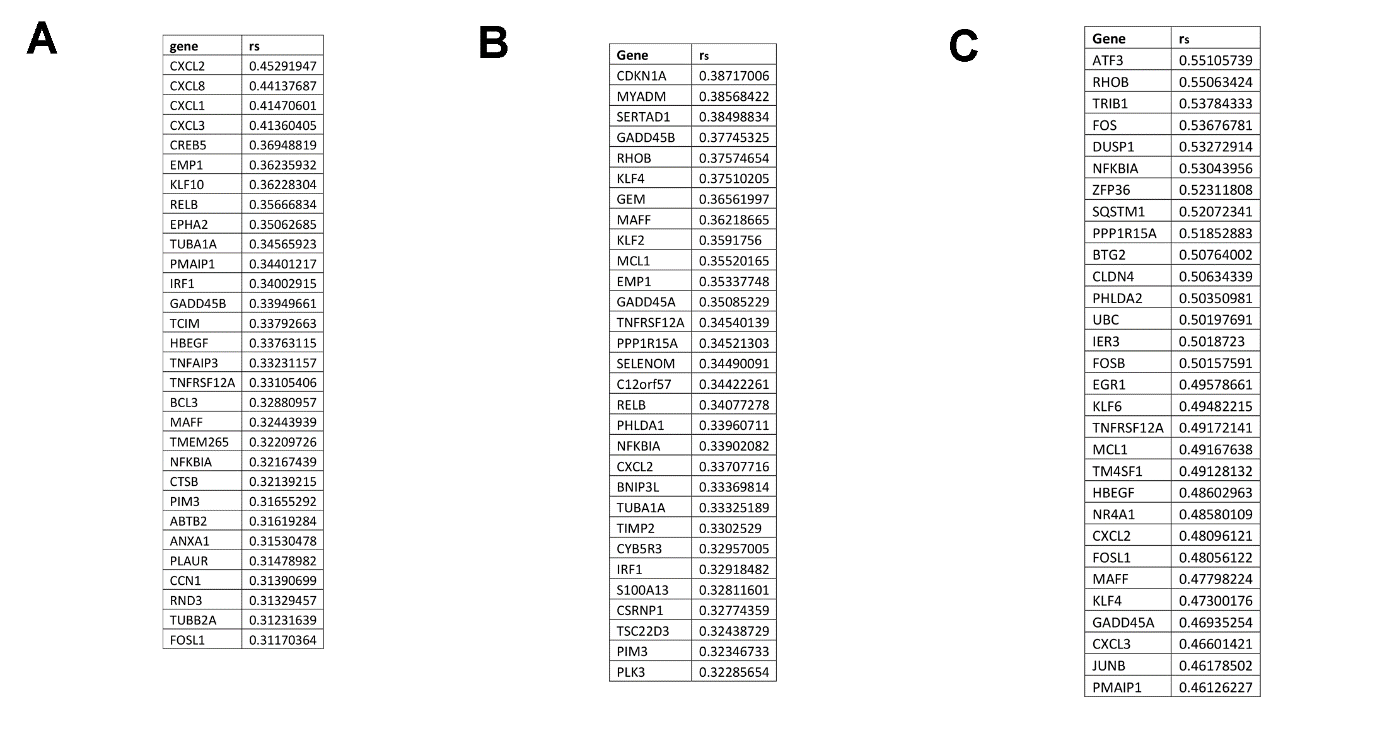

**Supplementary Table S4. Top 30 genes with positive correlation with HALLMARK_APOPTOSIS signature in treatment naïve (A), responders (B) and progressors (C), p < 0.001 for Spearman's rank correlation coefficient.**
