## Supplementary Fig. S4 for "Single-cell Transcriptome Profiling of Post-treatment and Treatment-naïve Colorectal Cancer: Insights into Putative Mechanisms of Chemoresistance"

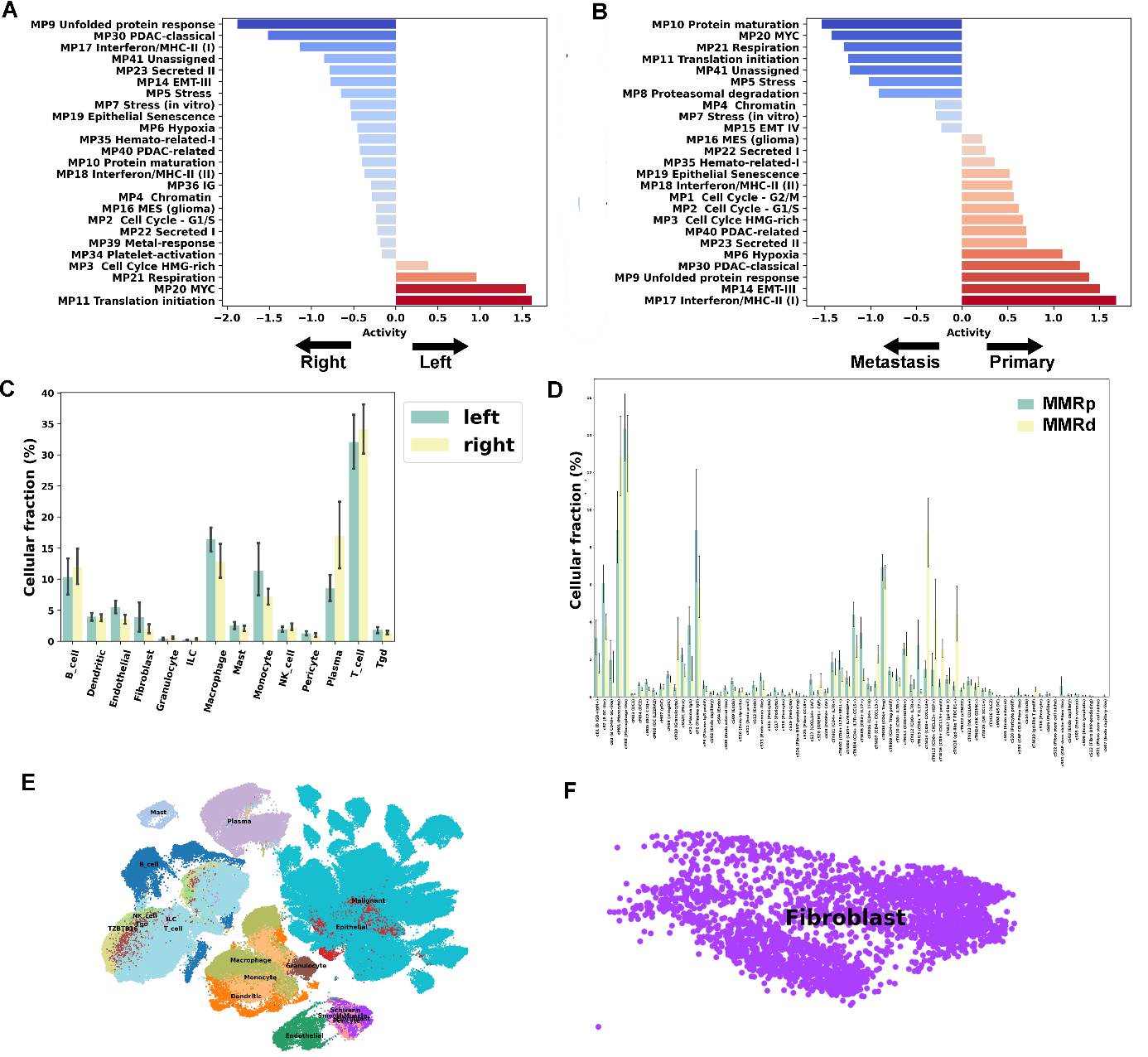

**Supplementary Figure S4. Analysis of validation treatment-naïve cohort (GSE178341).** Pathway-enrichment analysis for MSI compared to MSS tumors (A) and for left-sided compared to right-sided tumors (B). Cell type numbers alterations in MSI/MRRd versus MSS/MMRs tumors (C) and tumors left versus right (D). Uniform manifold approximation and projection (UMAP) embedding of 257,251 cells (E) and 3,031 cells of fibroblasts (F) after Leiden clustering.
