## Supplementary Fig. S4 for "Single-cell Transcriptome Profiling of Post-treatment and Treatment-naïve Colorectal Cancer: Insights into Putative Mechanisms of Chemoresistance"

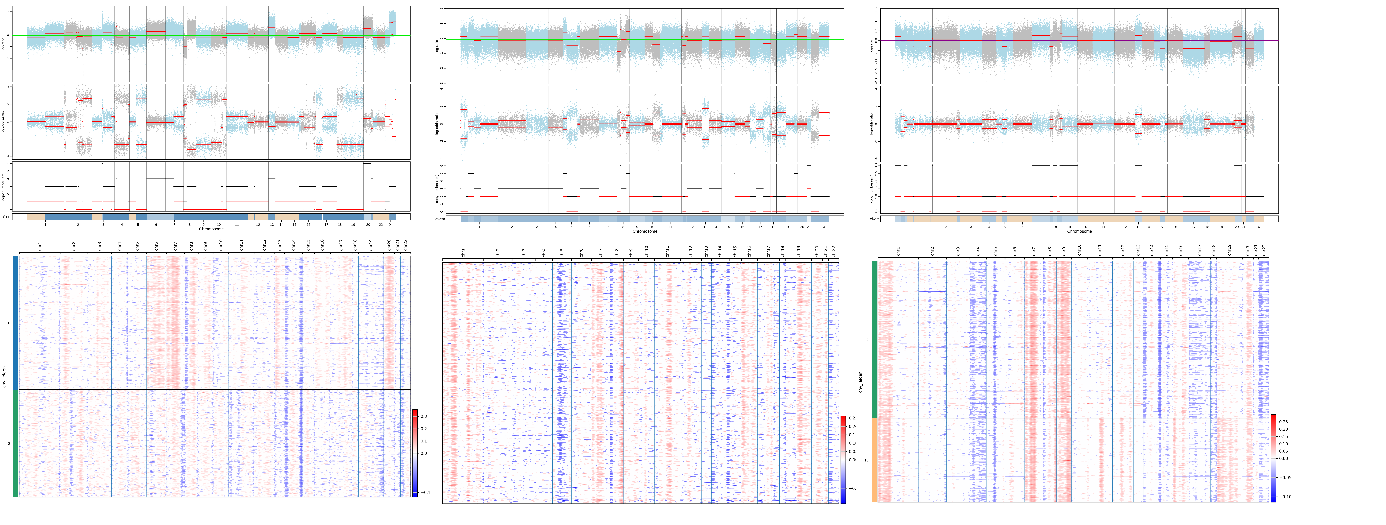

**Supplementary Figure S3. Comparison of CNAs in tumor-normal pairs from WES using FACETS and CNV heatmaps from scRNA-seq for 3 selected tumors.**
